# Comparison of localGEBV and Optimal Haplotype Stacking Fitness Functions using a Novel R Package: HapSelect

**DOI:** 10.64898/2026.07.08.737160

**Authors:** Will Shaffer, Victor Papin, Zane Carter, Stephanie Brunner, Jingyang Tong, Kira Villiers, Hannah Robinson, Kai P. Voss-Fels, Ben Hayes, Lee Hickey, Eric Dinglasan

## Abstract

Haplotype-based breeding strategies have emerged as promising approaches to maximize long-term genetic gain by identifying complementary parental combinations while maintaining genetic diversity. However, these methods typically require phased genotypes and more intensive workflow pipelines and skillsets. We developed a novel local genomic estimated breeding value (localGEBV) fitness function with similar intent to the optimal haplotype stacking (OHS) framework fitness function and implemented both in the novel R package, HapSelect. Our aim was to evaluate whether phased haplotypes provide additional benefit over the more easily available dosage-based unphased genotypes in highly inbred crops. A subset of bread wheat nested association mapping (NAM) population comprising 444 lines genotyped with 6,054 DArT-Seq markers was analysed. Marker effects were estimated using rrBLUP, localGEBV and haplotype effects were calculated across linkage disequilibrium-defined haploblocks, and genetic algorithms (GA) were used to identify optimal sets of 30 founders using either a localGEBV derived fitness function with unphased, dosage inputs or the OHS fitness function with phased inputs. Selected parental sets were compared with conventional truncation selection (TS) through 150 generations of forward simulation. The OHS fitness function achieved a marginally greater optimized ultimate GEBV than the localGEBV fitness function during GA optimization, with only 18 of the 30 selected founders overlapped between the two methods. Despite these differences, forward simulations demonstrated nearly identical long-term genetic gain for localGEBV and OHS-selected founders, with both approaches outperforming conventional truncation selection by maintaining greater genetic diversity and delaying the genetic plateau. The minimal difference between localGEBV and OHS is likely attributable to the high homozygosity of the population, where localGEBV and haplotype effects are nearly confounded. These results demonstrate that dosage-based localGEBV provides a practical alternative to phased haplotype approaches for parent selection in inbred crops, substantially simplifying genomic workflows while maintaining long-term breeding performance. Future work should evaluate these methods in more diverse inbred populations and outbred species, where great haplotypic diversity may increase the advantage of true haplotype-based optimizations.

## Introduction

Genetic technologies have been a key driver in rapid genetic improvement in the last century. Throughout most of human civilization phenotypic performance has been utilized to select genotypes for the next generation (Falconer et al., 1996; Bourdon, 2000). With the introduction of statistics, measurements were adjusted to be standardized across time, area, and other variables (Bourdon, 2000; Piepho et al., 2008). And towards the end of the last century, statistical design and pedigree-informed selection with selection indices or mixed model methodology enhanced selection accuracy further (Henderson, 1975; Bourdon, 2000; Meuwissen et al., 2001; Piepho et al., 2008). Most recently, genomic selection (GS) has been a key driver of genetic progress (Meuwissen et al., 2001; Piepho et al., 2008; VanRaden, 2008; Heffner et al., 2009). Each of these technological improvements have served to increase the rate of genetic gain by improving key aspects of the breeder’s equation, including selection accuracy, decrease the generation interval, and/or increase selection intensity. However, further tools are needed to further increase the rate of genetic progress and maintain useful genetic diversity, especially given the likelihood of climate change, disease pressure, and decreasing input (e.g., nitrogen) availability via government regulation or supply chain issues (Ray et al., 2015; Snowdon et al., 2021; Cooper and Messina, 2023; Quitzow et al., 2025). These forces all act in opposition within the breeder’s equation (Lush, 1937; Bourdon, 2000), decreasing selection accuracy through complex genotype-by-environment interactions, decreasing selection intensity if only a proportion of the population can thrive under these conditions (natural selection/fitness), increasing generation interval if crops and livestock face increased stress and time to reproductive maturity, and decreasing genetic variation through environmental pressures and intense natural selection.

Haplotype-based tools may contribute to the solution to increase the rate of genetic gain and maintain diversity. Haplotype methods have been gaining increasing popularity, with many proponents advocating for their use to increase selection accuracy and capture additional regional genetic effects. Other regional based genomic tools, such as local genomic estimated breeding values (localGEBV), which are GEBV at a given genomic segment rather than the whole genome, have also been proposed as an alternative to a true haplotype-based framework (Fan et al., 2011; Cole and VanRaden, 2011; Kemper et al., 2012; Voss-Fels et al., 2019; Shaffer et al., 2025). While both fundamentally rely on the same underlying theory to define regions and aggregate their effects, localGEBV utilizes the standard dosage-based genotypic framework to compute segmented GEBV with genomic prediction in comparison to allele-specific, true phased haplotype approaches (Kemper et al., 2012; Voss-Fels et al., 2019; Shaffer et al., 2025). If localGEBV can be utilized as an alternative, this would be advantageous because dosage-based single nucleotide polymorphisms (SNP) are more commonly available in crops and livestock than phased genotype data. An evolutionary genetic algorithm with an appropriate fitness function can then be utilized to select founders for mating and crossing, such as with the optimal haplotype stacking (OHS) fitness function and framework (Kemper et al., 2012; Villiers et al., 2024). The OHS framework attempts to identify individuals that maximize genetic potential while simultaneously requiring that haplotypes be derived from different founders to maintain genetic diversity. If all individuals from a population can be utilized, then the fitness function follows the concept of “the ultimate genotype” which aims to find the best combination of haplotypes under some series of restraints (Hayes et al., 2024). Other optimization algorithms (Yadav et al., 2025), or constraints, such as optimal haploid value (OHV; Daetwyler et al., 2015) and optimal population value (OPV; Goiffon et al., 2017), have been utilized in parent selection. In combination with different constraints, these genomic tools offer a more granular approach to optimize selection and mate allocation to maintain diversity and maximize genetic potential, similar to optimal contribution selection (Wray and Goddard, 1994), but at a local genomic level rather than consideration of whole-genome averages. Thus, short-term and long-term genetic progress may be improved or at least maintained (Villiers et al., 2024).

While a fitness function for true, phased haplotype approaches have been developed and characterized (Kemper et al., 2012; Villiers et al., 2024), it remains unclear whether unphased data can be utilized to realize the same benefits. OHS conventionally relies on phased haplotypes, whereas a localGEBV formulation can be used on standard dosage-based and unphased genotype calls. Phasing adds workflow complexity, technical expertise and labor cost, so establishing whether the localGEBV framework performs comparably on unphased data has practical breeding value. In highly inbred crops the case is particularly compelling. With high homozygosity, phased and unphased genotypes carry nearly equivalent information, meaning the two may identify similar founders and deliver comparable genetic gain. If so, breeders could rely on simple dosage-based data. We address this by (1) developing a localGEBV fitness function for unphased genotype data comparable to OHS, 2) comparing the founders selected by each approach using the same markers and inbred wheat population, and 3) implementing both in a novel R package, HapSelect, and using forward simulation to compare long-term genetic gain of OHS and localGEBV based parent selection using phased vs unphased haplotype datasets, respectively, along with the traditional truncation selection (TS) approach.

## Methodology

### Population

A subset of a bread wheat nested association mapping (NAM) population was analyzed in this study. The full dataset, comprising 1,819 lines phenotypes across 10 environments in Australia, was described in a previous study (Amin et al., 2025). Here, we analyzed grain yield (tonnes per hectare) from the subset of seven environments conducted at Warwick, Queensland, Australia, from 2014 to 2017. Each site contained anywhere from 162 to 975 lines with low to moderately high overlap (Table 1). Following Amin et al. (2025), plants were grown in 2 × 4 m plots (five rows at 25 cm spacing) with a target density of 100 plants m^−2^, and the full plot area (8 m^2^) was harvested for grain yield. One of the earliest trials (WAR14rf) used smaller 2 × 3 m plots to accommodate the larger number of genotypes.

**Table 1.** Number of unique genotypes in each environment (diagonal) and overlap between environments (off-diagonal).

| NAM11-WAR16RF | NAM12-WAR17RF | NAM13-WAR17IR | NAM3-WAR14RF | NAM4-WAR15RF | NAM5-WAR15IR | NAM6-WAR15RF |
| --- | --- | --- | --- | --- | --- | --- |
| 866 | 242 | 177 | 477 | 696 | 140 | 187 |
|  | 679 | 209 | 179 | 234 | 64 | 84 |
|  |  | 210 | 128 | 171 | 51 | 46 |
|  |  |  | 802 | 675 | 143 | 27 |
|  |  |  |  | 975 | 161 | 28 |
|  |  |  |  |  | 162 | 27 |
|  |  |  |  |  |  | 881 |

### Multi-Environment Trial

A multi-environment trial analysis was conducted in ASReml-R 4.2 (Butler et al., 2017) as per the following model:

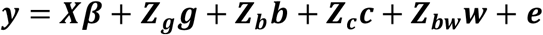

where ***y*** is the vector of yield observations for each genotype at the seven environments for reach replicate. The ***Xβ*** term contains the incidence matrix connecting fixed effects in ***β*** to observations in ***y*** and the fixed effects for the intercept and differential mean effects of each of the seven environments (set-to-zero restriction) to be estimated. ***Z***_***g***_ is the incidence matrix connecting each observation to the random genotype effects within a site and ***g*** is the vector of random genotype by environment effects, distributed as:

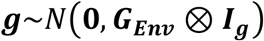

with ***G***_***Env***_ denoting the seven by seven genotypic covariance structure between environments and ***I***_***g***_ denoting the identity matrix (i.e., unrelated genotypes capturing broad-sense genotypic effects) with dimensions of the number of unique genotypes across all environments. The ***Z***_***b***_, ***Z***_***c***_, and ***Z***_***w***_ incidence matrices connect observations to block design, column, and row effects nested within environment and block design, column, and row effects have assumed distributions of the form:

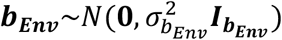

using block design as example. In this case, ***b***_***Env***_ is the vector of block design effects within an environment, assumed to have an environment-specific distribution and independence between block design effects both within and across environments. The same structure was utilized for rows and columns. That is, unique distributions of row and column effects were specified in each environment, recognizing independency between rows and columns within an environment, but also that row and column effects are derived from different distributions and are independent entities across environments. Finally, residuals were distributed:

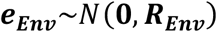

where ***R***_***Env***_ denotes the environment-specific residual covariance matrix. Within each environment, residuals were modelled using the direct sum of two first-order autoregressive spatial covariance structure across the row and column dimensions. Genotypic correlations across environments were then extracted. Broad-sense heritability, *H*^2^, was estimated using the ratio between the environment-specific genotype variance and sum of the environment-specific genotype variance and residual variance.

### Estimation of Overall Performance Best Linear Unbiased Estimates

Using correlated environments, the following model was fit in ASReml 4.2 to generate overall performance (OP) best linear unbiased estimate (BLUE) values for each genotype for yield:

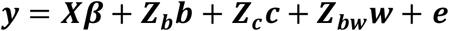

where all terms are as before, except for ***Xβ***, and records were only for 4 correlated environments. The fixed terms included the intercept, genotype effects, and environmental effects, with no interaction between the two because environments were correlated. Using the predict function in ASReml 4.2, BLUEs were computed as linear contrasts of the fixed terms to compute genotype main effects averaged over environment. Only the BLUEs for the Suntop NAM families, comprising 444 lines, were utilized in downstream genomic analyses to avoid excessive macro-scale population structure that evidenced in principle component analyses (PCA) when other NAM populations, like MACE, were included (data not shown).

### Genotypic Data

Genotypes of 18,827 DArT-seq unphased SNP markers were provided for all 444 NAM lines. Approximately 10,000 markers had known genomic coordinates after a basic alignment search tool (BLAST) search to align the marker sequence fragments to the Chinese Spring wheat genome assembly, often referred to as the CS-CAU genome assembly (Wang et al., 2025). Markers were then recoded into standard dosage format (0 = reference allele homozygote, 1 = heterozygote, and 2 = alternate allele homozygote) and subsequently filtered for a minor allele frequency less than 0.01, heterozygosity value greater than 0.15, call rate less than 0.80, and linkage disequilibrium (LD) threshold greater than 0.99. After filtering, a total of 6,054 markers remained. The LD values were generated using HapSelect (https://wrshf7.github.io/HapSelect-Docs/) as a wrapper to call PLINK 1.9 (Chang et al., 2015). Population structure was visualized utilizing PCA on the marker dosage matrix.

### Marker Effect Estimation and Cross-Validation Accuracy

Marker effects for yield OP were estimated using the 444 Suntop NAM OP BLUEs and 6,054 unphased, filtered dosage markers aligned to the CS-CAU genome assembly. HapSelect was utilized to estimate marker effects with a wrapper for the ridge regression Best Linear Unbiased Prediction (rrBLUP) R package (Endelman, 2011) with no addition effects outside of the marker random effects and the yield BLUEs as phenotypic input. The additive genetic variance was approximated by multiplying the restricted estimated maximum likelihood (REML) estimate of marker variance with the Van Raden method 1 (VanRaden, 2008) scaling term in the genomic relationship matrix denominator, ∑2*p*_*i*_*q*_*i*_ for marker *i*. Narrow-sense heritability of the OP was calculated in the same manner as the broad-sense heritability with this approximation of the additive genetic variance. HapSelect and rrBLUP were further utilized to predict cross-validation (CV) accuracy, defined as the correlation between genomic estimated breeding value (GEBV) and BLUEs using a bootstrapping resampling technique. Thirty iterations of CV were conducted with 80% of the data sampled and 20% masked for validation.

### Phased Marker Matrix Construction

In addition to the unphased standard dosage (allele count) matrix, a phased marker matrix was constructed to compare a true, phased haplotype approach to localGEBV in parent selection. Given the NAM lines were relatively inbred (i.e., F_4_-derived) with an expectation of 12.5% of the original heterozygosity remaining, the heterozygous genotype calls were randomly phased to the chromosome. This level of inbreeding was considered appropriate given that many plant breeders who work with inbred crops often bulk lines from the F_4_ stage onwards and treat them as effectively fixed.

The use of Beagle to properly phase will be explored in the future.

### Haploblocking

Haploblocking was conducted using an LD threshold of 0.3 and tolerance of 3 as described in the HapSelect documentation and various studies (Voss-Fels et al., 2019; Shaffer et al., 2025), and implemented in the HapSelect package; however, these thresholds should be tailored to the specific population under investigation. Briefly, the algorithm starts at the first SNP in the chromosome and compares the LD to the next SNP. If the LD between the two SNP meets the threshold, the SNP are grouped into a haploblock and the algorithm will then compare the newly added SNP to the next adjacent SNP. This process continues until a SNP pair fails to meet the LD threshold. If the LD between the two SNP under consideration does not meet the threshold, then the tolerance is incremented by one and the LD between the last successful SNP and the next SNP after the previously failed SNP is compared. If multiple SNP in a row (enough to surpass the max tolerance count) fail to meet the threshold, then the block is terminated and all failed SNP are not added to the block. If the LD between the last successful SNP and the next SNP under evaluation meets the threshold, then the new SNP and all failed SNP in between are added to the haploblock and the tolerance counter is reset. The purpose of the tolerance parameter is to accommodate issues with marker quality (such as genotyping bias, incorrect genotype calls, or missing calls biased to certain genotypes) and marker positions based on a single reference genome that may affect LD and to accommodate misplaced markers (Shaffer et al., 2025). The summary information regarding haploblock size and marker content is presented in Table 2.

**Table 2.** Haploblock size distribution for a linkage disequilibrium threshold of 0.3 and tolerance of 3, including the mean number of single nucleotide polymorphisms (SNP) per haploblock for non-singleton haploblocks, the max number of SNP in a haploblock, mean base pairs (BP) in megabases (Mb) for non-singletons, max BP, number of singleton (singleSNP) haploblocks, and proportion of singletons.

| Mean SNP | Max SNP | Mean BP (Mb) | Max BP (Mb) | Singletons | % Singleton |
| --- | --- | --- | --- | --- | --- |
| 5.71 | 112 | 13.93 | 455.81 Mb | 417 | 39.3% |

### Calculation of localGEBV, Haplotype Effects, and Haploblock Variance

Calculation of localGEBV and haplotype effects follows the same paradigm as genomic selection. In genomic selection, 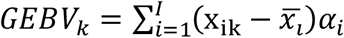 where the GEBV of individual *k* is the sum of all of individual *k*’s centered (2*p* as the centering value in a diploid) dosage genotypes (VanRaden, 2008) across *I* markers multiplied by the estimated marker (allele substitution) effect previously described from rrBLUP. This centering prevents biasing GEBV and is an important step. Similarly, consider an individual haploblock, block *j* of *J* total haploblocks, and the corresponding markers in that block, *i* ∈ *j*. Then, 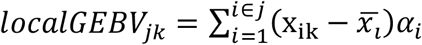 (Voss-Fels et al., 2019; Shaffer et al., 2025). That is, the localGEBV for an individual is the sum of the centered, unphased genotypes multiplied by the marker (allele substitution) effects corresponding to those centered genotypes in a given block. It then follows that the sum of localGEBV is equivalent to the GEBV for an individual: 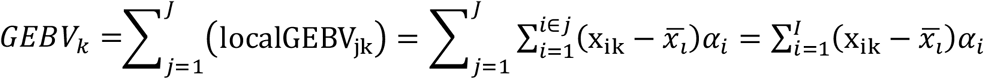. Thus, expectations and variance of whole-genome GEBV and individual loci can be similarly applied to localGEBV (e.g., the expectation of block *j* is 0).

Haplotype effects were similarly calculated, but on a per chromosome basis for each of the two chromosomes (diploid) at each block with phased data: 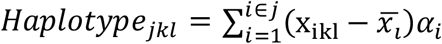 where terms are as before, but *Haplotype*_*jkl*_ is the additive haplotype effect of individual *k* at block *j* on chromosome *l* ∈ {1,2} (Kemper et al., 2012; Villiers et al., 2024). It follows for a diploid that 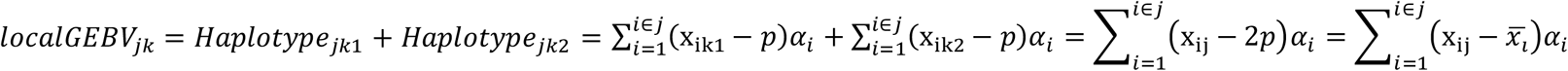.

Finally, the variance at haploblock *j* was considered analogous to the variance of whole-genome GEBV in the decomposition of genetic variance into variance captured by the data versus prediction error variance (Henderson, 1975), and was calculated as:

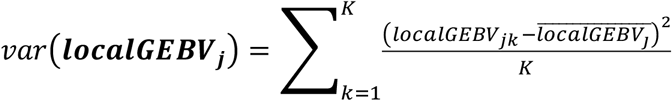. Importantly, this represents the variance of the estimates or the variance of the total genetic variance at block *j* that could be captured with the data and *K* is the total number of individuals (444).

### Selection of Haploblocks and Parent Selection with Fitness Functions and Genetic Algorithms

Haploblocks were arranged in descending order by localGEBV haploblock variance. The criteria utilized for block selection was the number of haploblocks that explained at least 95% of the cumulative haploblock variance, which was 246 of 1,060 total haploblocks. For parent selection and simulation, the objective was to select 30 individuals from the 444 individuals in the Suntop NAM population that would be most synergistic and could be utilized to create the best “ultimate genotype” (Hayes et al., 2024) under the constraint of utilizing only 30 individuals. This is a highly combinatorial problem with: 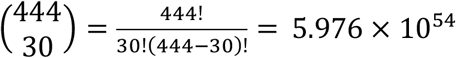 unique combinations of individuals. This number of computations becomes unfeasible when trying to compare the ultimate GEBV across each combination. Thus, a genetic algorithm (GA) heuristic optimization framework using the GA R package (Scrucca, 2016) implemented in HapSelect was utilized to find an optimized combination using a custom fitness function for localGEBV and the OHS fitness function (Kemper et al., 2012; Villiers et al., 2024).

The localGEBV fitness function was defined as follows for a given combination (set of 30 individuals): 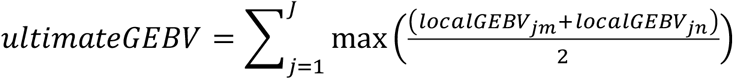, where *localGEBV*_*jm*_ and *localGEBV*_*jn*_ are the localGEBV at block *j* of individuals *m* and *n* in the parental set (*m* ∈ {1 … 30} and *n* ∈ {1 … 30}) and 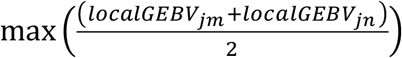 represents the best average localGEBV of two individuals out of 30 possible founders at block *j*. The constraint of *m* ≠ *n* was added to prevent selfing being part of the optimization criteria and make the outcome more comparable to the OHS framework. Furthermore, the value of *m* and *n* could vary across each block *j*, such that the best segments were chosen for each block. Thus, the ultimate GEBV of a given combination was the sum of the best possible localGEBV average from all 30 founders. This fitness function was utilized with the GA optimization framework from the GA package in HapSelect.

The OHS haplotype fitness function was specified in much the same manner, except each individual contained two haplotypes (diploid) and the OHS framework has the constraint that haplotypes must come from two individuals (Kemper et al., 2012; Villiers et al., 2024). The fitness function was specified as follows: 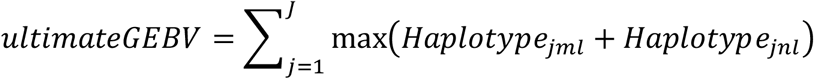. Where terms are as before and *m* ≠ *n*. That is, the two haplotypes must be derived from two different founders. Importantly, the two haplotypes could be identical by state (i.e., have identical allele sequences), provided they were inherited from different founders (Villiers et al., 2024).

The fitness functions were implemented with the GA algorithm from the GA package in HapSelect. For founder selection, the number of founders to select was specified as 30, with 100 parent pools per cycle, a mutational rate of 0.5, a crossover rate of 0.5, and a maximum number of iterations of 3000. The mutational parameter and crossover parameter specified gave each parent pool a 50% chance to randomly swap one individual for a different individual from the entire population not in the parent pool and for a pair of parent pools to swap half their members. If there were non-unique individuals after swapping, the duplicated individuals were mutated with a unique, random individual from the total population similar to the mutation parameter. At the end of the optimization, the best set of 30 was utilized for downstream simulation.

### Forward Simulation

Simulation was conducted with the GenomicSimulation package (Villiers et al., 2022) implemented with a wrapper in HapSelect. For simulation, the whole-genome marker data and marker effects were utilized for the 30 localGEBV-selected founders, the 30 OHS-selected founders, and the 30 best whole genome GEBV founders, denoted as the TS-selected founders. For the TS and localGEBV-selected founders, the heterozygous genotypes were randomly phased, similar to the haplotype marker matrix. A genetic map was not available, so the genetic map position was assumed to be proportional to the marker position on the chromosome with a max of 100 centiMorgans (cM) per chromosome. That is, the genetic map position, in cM, of marker *i* was: 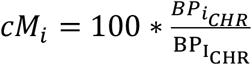, where *cM*_*i*_ is the map position of marker *i*, 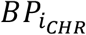 is the base pair position of marker *i* on its constituent chromosome, and 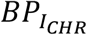 is the base pair position of the last marker on a given chromosome. For simulation, recurrent TS was applied to the localGEBV, OHS, and TS selected parent sets for 150 generations. Each generation 200 progeny were generated with random mating using all possible crosses, and 30 genotypes were selected with TS to proceed to the next generation.

## Results and Discussion

### Multi-Environment Trial Analysis

The distributions of raw yield phenotypes in each of the seven environments and broad-sense heritabilities are presented in Supplementary Fig. 1. The raw yield mean and variance between the NAM4-WAR15RF (rain-fed in 2015), NAM5-WAR15IR (irrigated in 2015), and NAM13-WAR17IR (irrigated in 2017) were markedly similar. However, the broad-sense heritability was notably higher in NAM13-WAR17IR than the other two. Other environments had greater variability, but also greater broad-sense heritability, such as NAM6-WAR15RF. However, there did not appear to be any overall trend between broad-sense heritability and raw, phenotypic variability within an environment or between broad-sense heritability and the environmental mean. There also did not appear to be any broad relationships between year and broad-sense heritability or irrigated versus rain-fed environments and broad-sense heritability. But, it should be noted this wasn’t explicitly tested or of primary interest in this particular study. Interestingly, we realized greater broad-sense heritability estimates than the original study characterizing this dataset (Amin et al., 2025). But, this could be caused by differences between a fully unstructured model versus a more parsimonious factor analytics (FA) model.

The genetic correlations between yield across environments were positively correlated but varied across the environments (Supplementary Fig. 2). Results suggested grouping all three 2015 environments and the NAM13-WAR17IR would be appropriate to generate overall performance (OP) BLUEs. These four environments were selected to generate a single OP BLUE per genotype that provided a robust phenotypic basis for estimating the marker and haplotype effects underpinning the parent-selection comparison, with their consistently moderate-to-high genetic correlations indicating they capture a common genetic signal suitable for combination. However, in contrast to the original study on this dataset, we realized greater genetic correlations between environments (Amin et al., 2025). This is also likely because of differences between the unstructured and FA model. Even so, all of the environments were also clustered into the same envirotype in Amin et al. (2025), further indicating the choice of these environments to generate OP BLUEs is appropriate.

### Overall Performance Best Linear Unbiased Estimates, Genotypic Data, Marker Effects, Haploblocking, and Haploblock Variance

The OP BLUEs were modeled and then the 444 Suntop NAM family lines were specifically extracted for extended analyses. The distribution of yield BLUEs for the 444 Suntop NAM lines are presented in Supplementary Fig. 3. The mean yield OP BLUE was 6.08 *t ha*^−1^, with a variance of 0.28 (*t ha*^−1^)^2^. This was comparable to the observed yield amongst genotypes in envirotype 1 in Amin et al. (2025) and to wheat yields in general in Australia (Hunt et al., 2019).

The distribution of markers pruned to these 444 lines is presented in Supplementary Fig. 4A. In general, the marker density was greatest in telomeric regions than in centromeric regions. While seemingly problematic, LD between markers is generally low in telomeric regions and high in centromeric regions because recombination is generally biased towards telomeres (Lloyd, 2023). Thus, far fewer markers are needed in centromeric regions to adequately capture effects because of extensive LD between markers and quantitative trait loci (QTL) allowing for efficient capture of QTL effects with far fewer markers (Solberg et al., 2008; Habier et al., 2009; Liu et al., 2015). This was supported by LD patterns because LD was quite extensive in this population (Supplementary Fig. 4B), especially in centromeric regions (Fig. 1A). Average genome decay across telomeric and centromeric regions was greater than 10 Mb to reach *R*^2^ = 0.20. Extensive LD would be expected in NAM lines given the family structure where donor parents were crossed to the elite parent Suntop.

**Figure 1.**
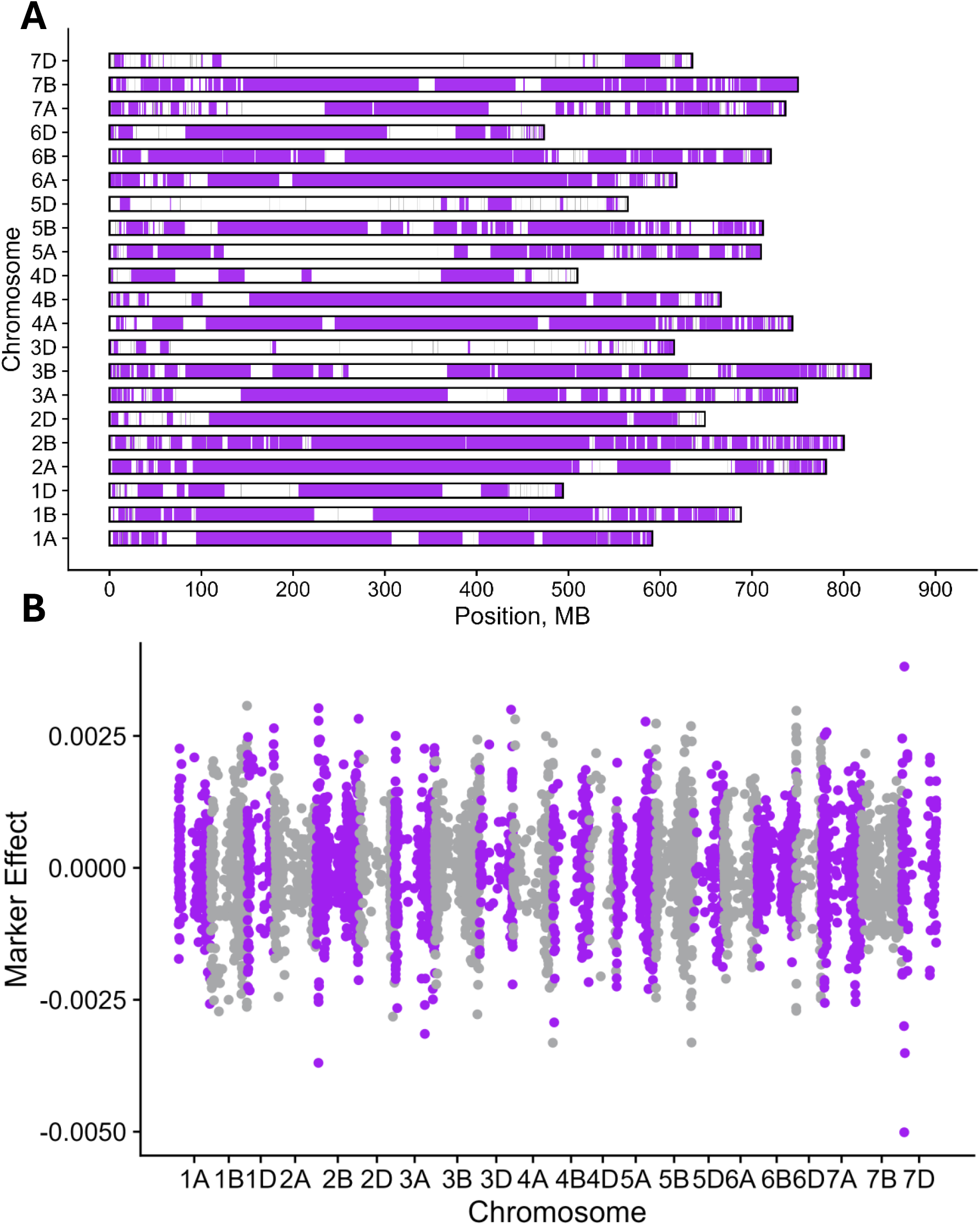
A. Distribution of haploblocks across the 21 wheat chromosomes for linkage disequilibrium threshold 0.3, tolerance 3, and positions in megabases (Mb). B. Manhattan plot of marker effects from ridge regression best linear unbiased prediction across the genome (rrBLUP).

Marker effects were then estimated with rrBLUP using the pruned, unphased dosage genotype matrix for the 444 NAM Suntop lines and OP BLUEs. The narrow-sense heritability was approximated at 0.30 with a mean CV prediction accuracy across 30 bootstrapped replicates of 0.44 and the marker effects are presented in Fig. 1B. While one marker stood out on chromosome 7D and a few intermittently on other chromosomes, marker effects were generally infinitesimally distributed, which is expected given the assumptions of rrBLUP (Endelman, 2011) and heavy shrinkage applied to large variance markers. There was a general bias towards marker effects in telomeric regions being greater, however. This may be because of the extensive blocking structure observed in the centromeric regions (Fig 4A). In such a case, markers would become highly collinear amongst each other and likely with QTL in the region and effects and variance would be split amongst the markers, which has been observed previously (Shaffer et al., 2025). In contrast, narrow blocking regions, such as the telomeres, would require greater marker density, but some markers would essentially capture the majority of a single QTL effect.

In support of this conclusion, large localGEBV effects (Fig. 2A) and large haploblock variances (Fig. 2B) were more uniform between telomeric and centromeric regions. Thus, the phenomena of localGEBV to aggregate dispersed marker effects described in Shaffer et al. (2025) was also demonstrated in this population. The unique, phased haplotype effect Manhattan plot, however, was not presented because it was nearly identical (except half the effect size) to the unique, unphased localGEBV Manhattan plot in Fig. 2A. This is likely because of the low heterozygosity in the genome indicating that phased haplotype effects were nearly perfectly correlated with unphased localGEBV values (i.e., 0.5 the localGEBV value given homozygosity causing a near double haploid state). In summary, there was very little overlap between large marker effects and major haploblocks (defined as high variance haploblocks). This demonstrates the power of the unphased localGEBV and phased haplotype methods to both aggregate dispersed effects in high LD regions in a genomic prediction framework.

**Figure 2.**
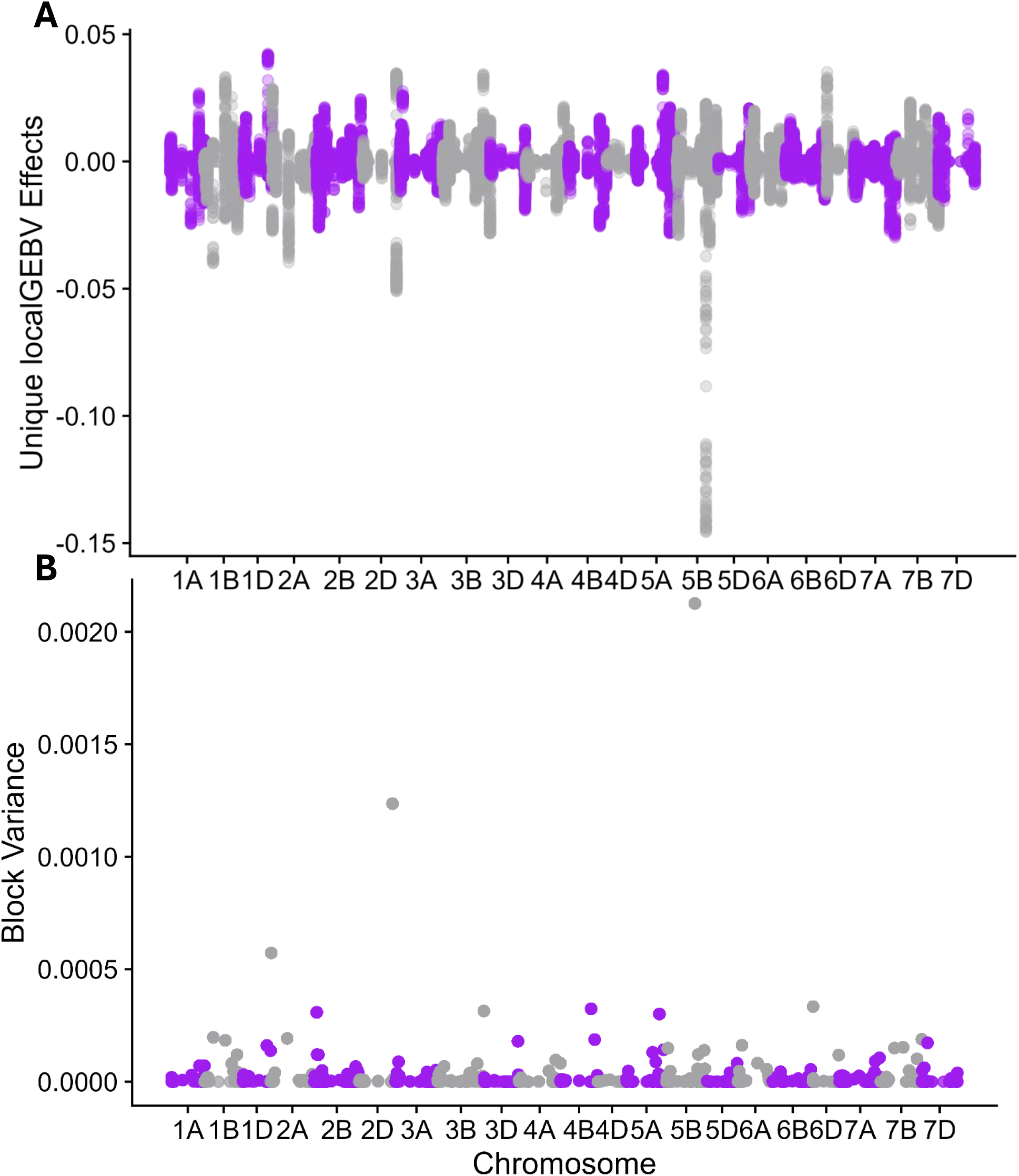
A. Distribution of unique, unphased local genomic estimated breeding values (localGEBV) effects across the genome. B. Distribution of haploblock variances across the genome based on localGEBV at a given block.

### Haploblock Selection, Genetic Algorithm, and Fitness Functions

Selection of haploblocks to use as input into the GA and fitness functions was based on the unphased localGEBV haploblock variance. A minimum threshold of blocks sorted in descending order based on haploblock variance explaining at least 95% of the cumulative haploblock variance was utilized to select blocks. Out of the 1060 blocks, 246 explained at least 95% of the cumulative variation. This number decreased substantially as the threshold was decreased from 95% to 90% or lower (data not shown). Selecting a subset of blocks is highly advantageous, because the vast majority of genetic variation is captured and the number of computation units is vastly decreased. This led to drastically decreased runtimes in the GA, because the number of computations per population in each iteration was decreased by nearly 77%. While optimizing based on the whole genome is better to maximize accuracy of selection, a trade-off may be needed for GA runtime performance if marker sets or populations grow substantially larger in practical breeding scenarios.

Using the 246 haploblocks, the maximum ultimate GEBV of the 100 parent pools across GA iterations for the phased haplotype and unphased localGEBV methods were computed and are presented in Fig. 3A. As might be expected, the phased haplotype-based approach achieved a slightly greater value than the unphased approach. This is expected, because the phased haplotype fitness function is more granular and dissects individual chromosomes compared to the unphased localGEBV fitness function. The localGEBV fitness function, in contrast, must average over chromosomal sets within an individual because of the unphased dosage input format. Thus, unless phased haplotypes are completely identical on both chromosomes within an individual, the unphased localGEBV will lose some power compared to the phased haplotype approach. The homozygosity and thus low heterozygosity across the population, which gives limited haplotype diversity, likely explains the relatively low difference between the parent sets, despite an overlap of only 18 founders between the OHS and localGEBV identified founders. In an outcrossed species with high heterozygosity, however, the unphased haplotype method may pull even further ahead because of the averaging problem. Evaluation in livestock and hybrid crops remains a future endeavor to assess functional differences between the two fitness functions.

**Figure 3.**
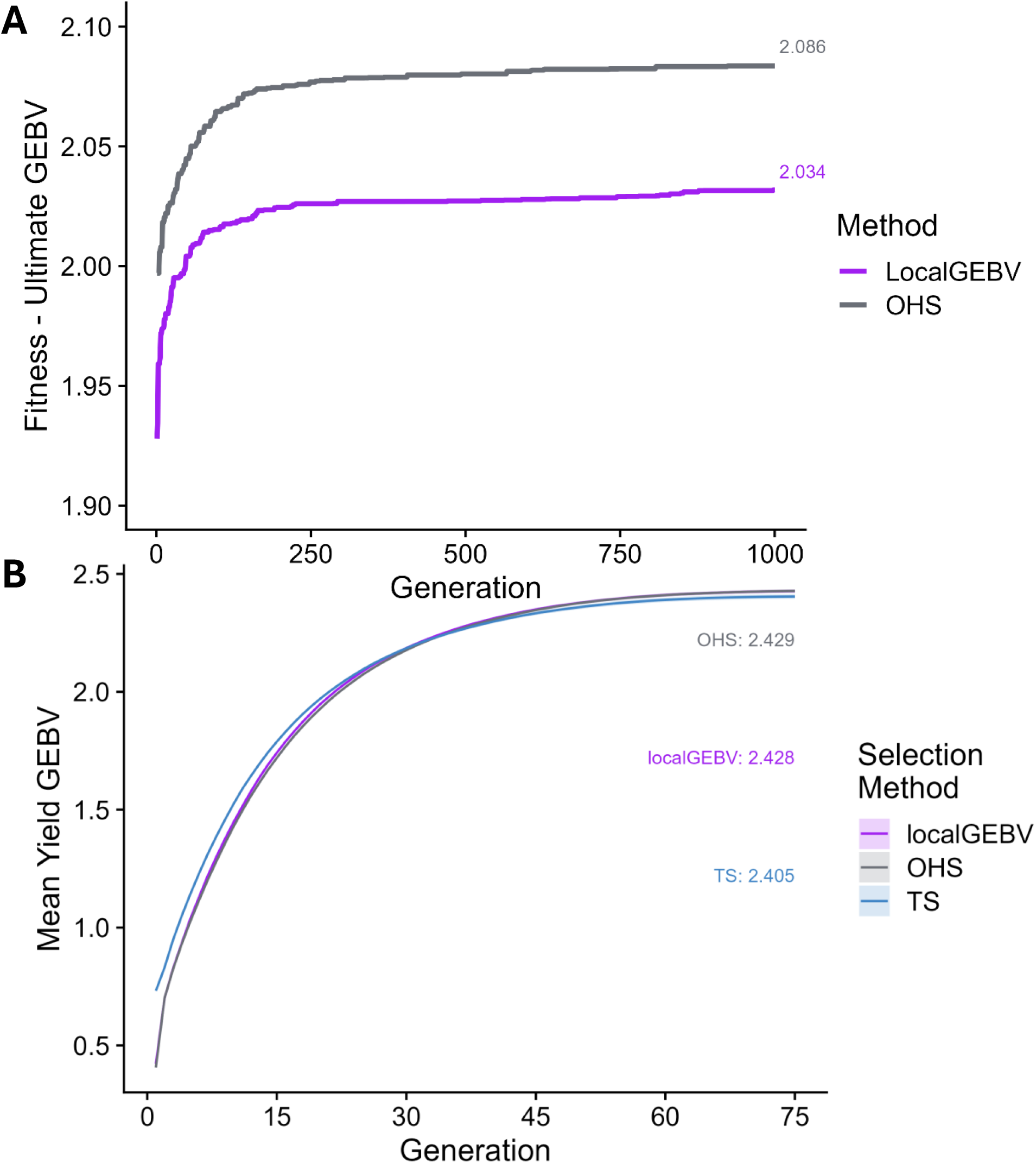
A. Maximum ultimate genomic estimated breeding value (GEBV) using a localGEBV fitness function for unphased, dosage inputs in a genetic algorithm (GA) vs an optimal haplotype stacking (OHS) fitness function for phased inputs in the GA. A total of 3000 iterations were run, but only the first 1000 are shown for clarity along with the maximum at 3000 displayed. B. Recurrent truncation selection (TS) simulation on founders derived from the localGEBV fitness vs OHS fitness functions in the GA and TS-selected founders using genomicSimulation. The mean value for each method of selection in the final generation was reported numerically in the plot along with +/- 1 standard deviation of replicates as a transparent ribbon. Only the first 75 of 150 generations are shown for clarity, but the maximum at 150 is displayed.

### Forward Simulation and Genetic Potential

Following selection of the 30 founders by each approach (localGEBV, OHS, and TS), forward recurrent TS was applied to each set, with results presented in Fig. 3B. As might be expected, the starting value of TS selected founders was higher initially (approximately 0.30 tonnes per hectare greater). This is because the GA does not prioritize a high overall mean GEBV, but instead prioritizes either unphased dosage or true phased haplotype complementarity. Thus, an individual can be chosen if it makes a useful contribution despite having a lower overall GEBV. This is how the GA prioritizes maintaining diversity for long-term genetic gain. As can be seen in Fig. 3B, both the localGEBV and OHS (haplotype) selected founders surpassed the TS-selected founders long-term despite having substantially lower initial mean GEBV, mirroring the results of (Villiers et al., 2024). It was surprising there was little difference in average GEBV after many recurrent TS cycles between TS-selected founders and localGEBV and OHS-selected founders, unlike in previous studies (Villiers et al., 2024; Tong et al., 2025). This may be because the NAM population was developed using a limited number of parental lines and may have contained limited haplotype diversity evidenced by high relationships in the genomic relationship matrix (Supplementary Fig. 5), which would cause the results to be nearly identical despite the apparent diversity in Fig. 4.

**Figure 4.**
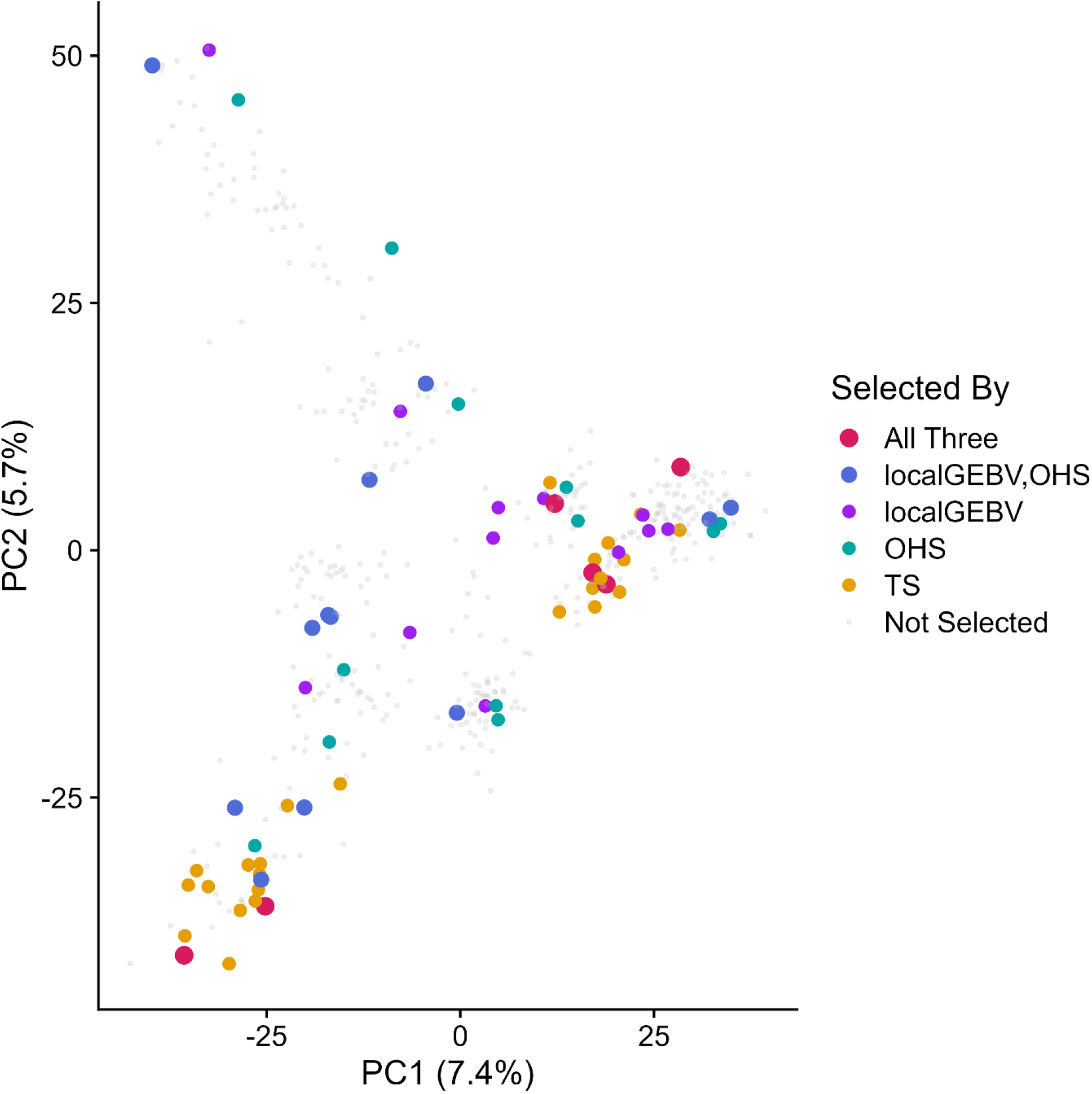
Principal component analysis (PCA) plot of the whole-genome marker data. Individuals are sized and colored by which method of selection chose them as founders: local genomic estimated breeding value (localGEBV) fitness function using unphased dosage inputs in the genetic algorithm (GA), optimal haplotype stacking (OHS) fitness function using phased inputs in the GA, truncation selection (TS) of whole genome GEBV, some overlap of the methods, or not selected by any combination of methods. Dot size represents number of methods that selected each individual.

In addition, localGEBV and OHS founder sets plateaued later on, indicating the localGEBV and OHS founders contained greater initial genetic variation than the TS-selected founders. This was apparent in the PCA structure in Fig. 4, where TS-selected founders clearly clustered to two subsets of the plot whereas OHS and localGEBV founders were sourced more uniformly from the entire population. Interestingly, TS-selected founders that overlapped with localGEBV or OHS GA-selected founders were only in common to all three methods, but not to TS-selected founders and localGEBV founders or TS-selected founders and OHS founders. In summary, the design intention of the two local genomic (localGEBV unphased dosage and phased, true haplotype) methods to select diverse, but complementary individuals and the increased diversity evidenced from the PCA was verified by long-term recurrent TS, because the simulations indicated greater genetic variation in the initial OHS and localGEBV founders.

Interestingly, despite a small, but meaningful difference between the localGEBV and OHS GA performance, the recurrent TS simulation showed little to no difference in long-term performance between OHS and localGEBV parent selection methods, even with an overlap of only 18 founders between the two local genomic methods. This may be because of the inbred nature of wheat, which would cause unphased, dosage localGEBV and phased haplotype effects to be nearly perfectly confounded. In a perfectly homozygous individual, the localGEBV are simply 2 times an individual haplotype effect (e.g., a double haploid) and would be perfectly confounded in parent selection of the localGEBV and OHS fitness functions. Both methods, however, should be compared in an outcrossed species, like livestock, to determine if there are functional differences between the two methods. Inclusion of a proper genetic map would also be informative to determine if recombination hotspots may influence long-term genetic potential in simulation. However, for inbred crops, there appears to be little to no functional difference between OHS and localGEBV fitness functionsin long-term population performance. Assuming this difference was not underestimated by the NAM structure, this is great news for breeders utilizing inbred species because it vastly simplifies the genomic workflow and input costs if simple SNP dosage genotype calls can be utilized for equivalent performance.

## Conclusions

A localGEBV fitness function (unphased, dosage inputs), derived from combining concepts from whole genome genomic prediction, was evaluated against the OHS (phased inputs) fitness function in an inbred wheat NAM population. Both methods showed superiority in maintaining diversity and long-term genetic compared to standard TS, but there was little to no difference in long-term potential between the localGEBV and OHS selected founders, despite a small difference in the GA. This was almost assuredly a consequence of the inbred nature of the crop, where unphased, dosage-based localGEBV are nearly perfectly confounded with phased additive haplotype effects. However, the performance of the two may differ in outbred species like livestock or hybrids and will be a future area of research. The small difference between OHS and localGEBV may also partly be a consequence of the NAM population structure and may be more pronounced in a population with more diversity. The lack of difference in this study is a boon to breeders, however, because it allows the usage of unphased dosage genotype calls and vastly simplifies the workflow, computation burden, and likely cost of use compared to OHS using phased genomic data in inbred species.

## Acknowledgements

ED was supported by the Queensland Government’s Advance Queensland Industry Research Fellowship Program (AQIRF096-2023RD6). LTH was supported through an ARC Future Fellowship (FT220100350). WS, VP and JT were supported as Adjunct Postdocs of the Australian Research Council Training Centre in Predictive Breeding for Agricultural Futures (IC230100016). The authors also thank Dr Jack Christopher and the project team for developing the wheat nested association mapping population, funded by the Grains Research and Development Corporation of Australia (Project UQ00068).

---

**Supplementary Figure 1.**
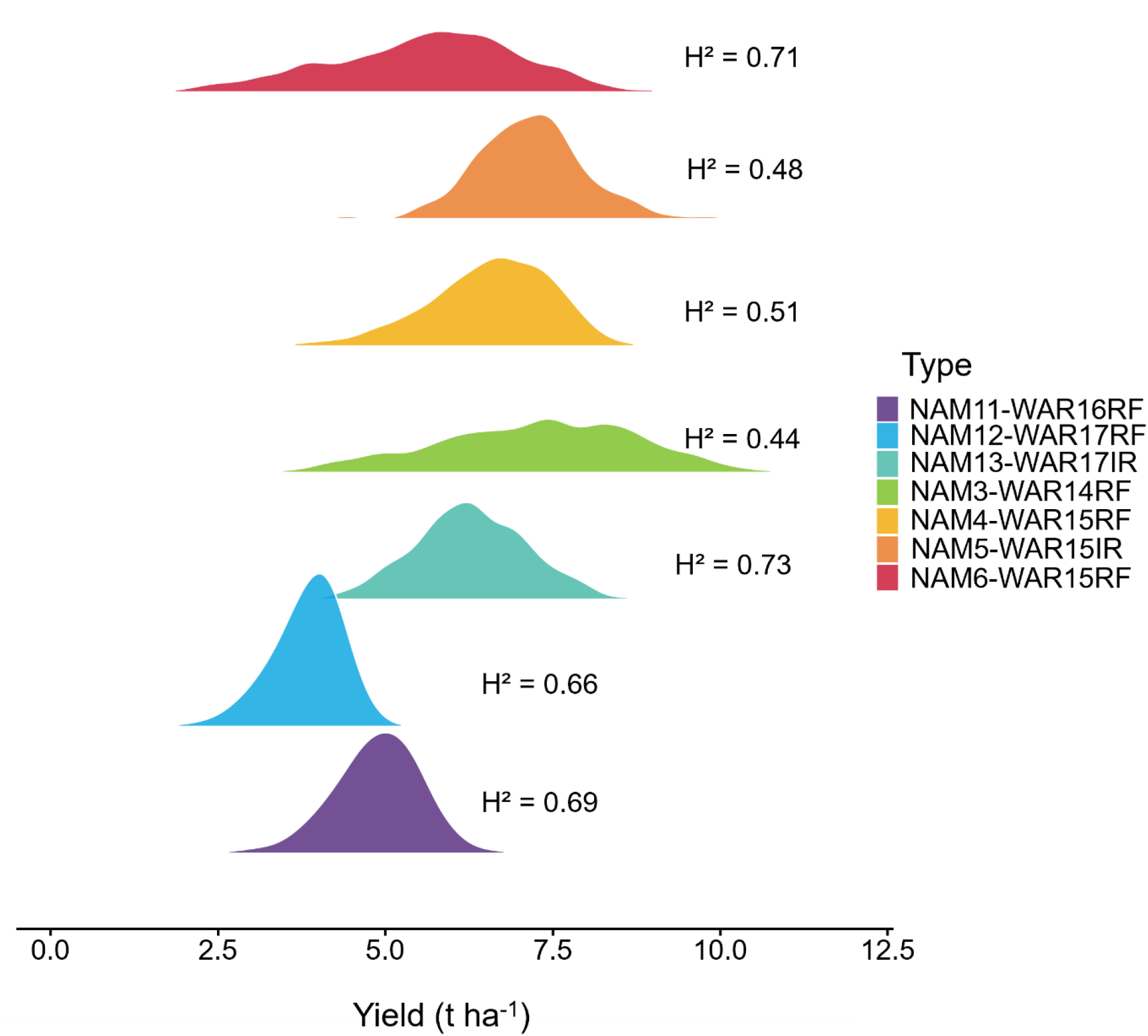
Density ridgeplots of yield in tons per hectare (*t ha*^−1^) in the seven environments (colors) evaluated with phenotypic observations as dot points. Broad-sense heritabilities were estimated from the multi-environment trial analysis.

**Supplementary Figure 2.**
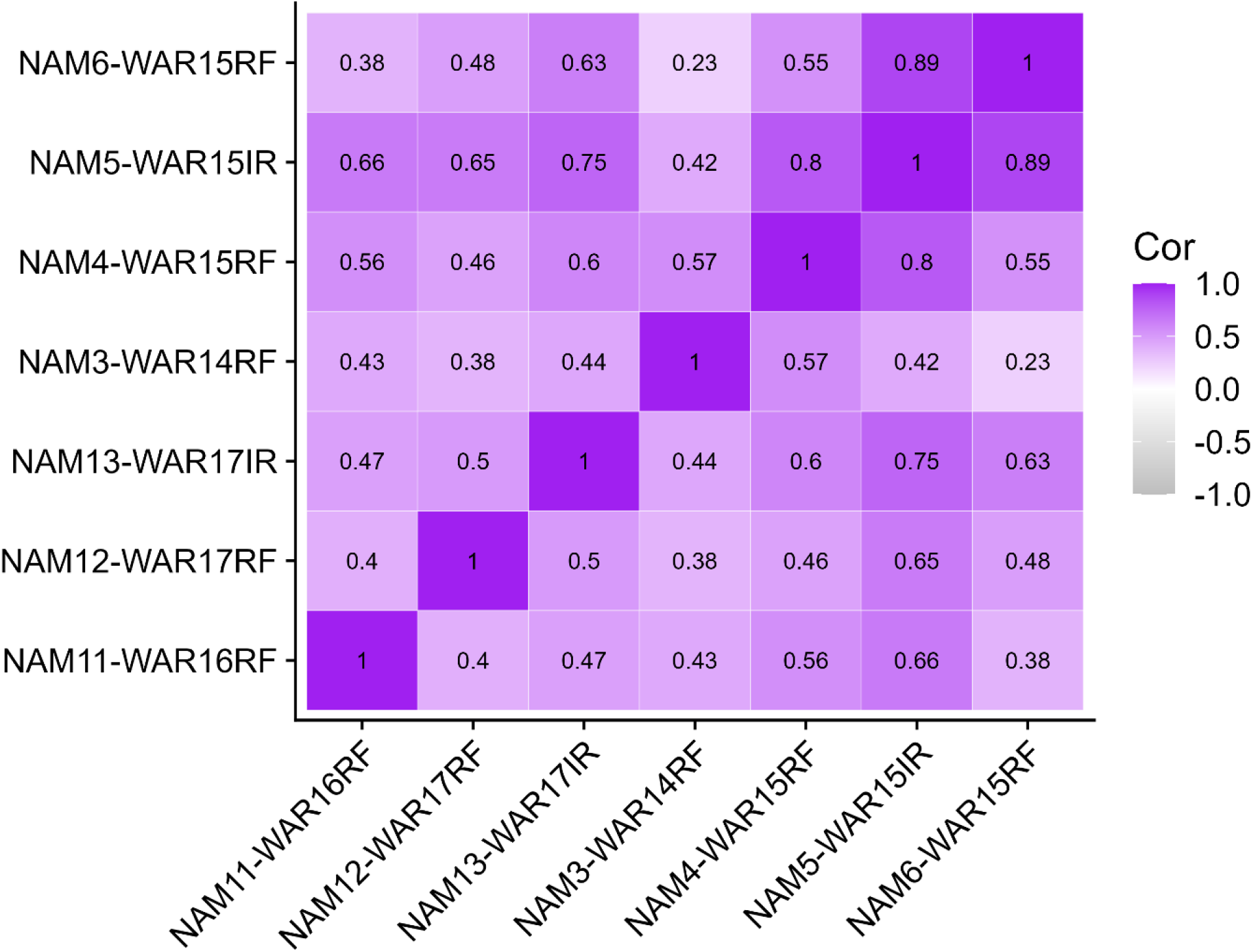
Multi-environment trial (MET) genotypic correlations for grain yield across the seven environments.

**Supplementary Figure 3.**
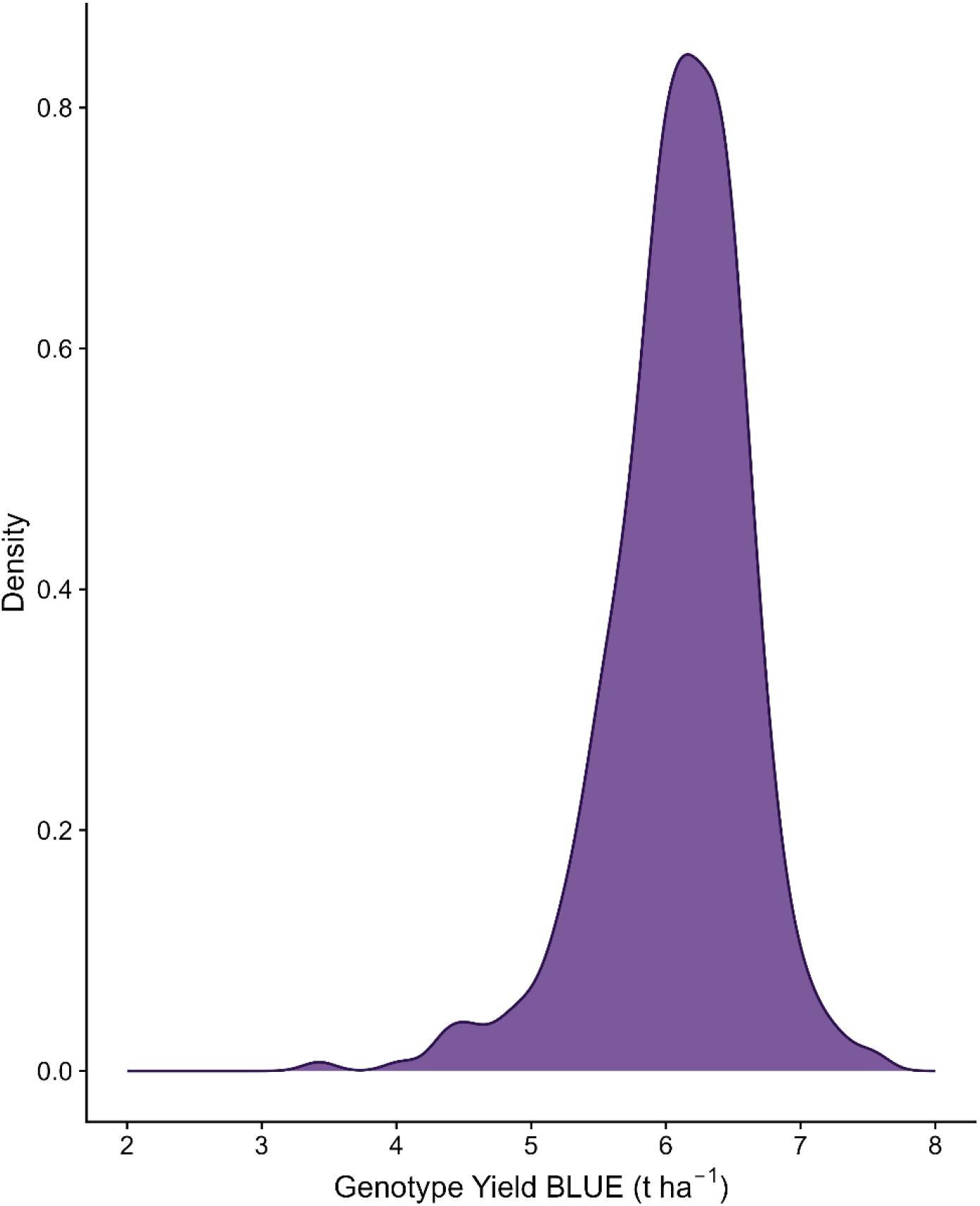
Distribution of best linear unbiased estimates (BLUE) for yield (tons per hectare, *t ha*^−1^) of the 444 nested association mapping (NAM) Suntop lines evaluated across four correlated environments (NAM6-WAR15RF, NAM5-WAR15IR, NAM4-WAR15RF, and NAM13-WAR17IR).

**Supplementary Figure 4.**
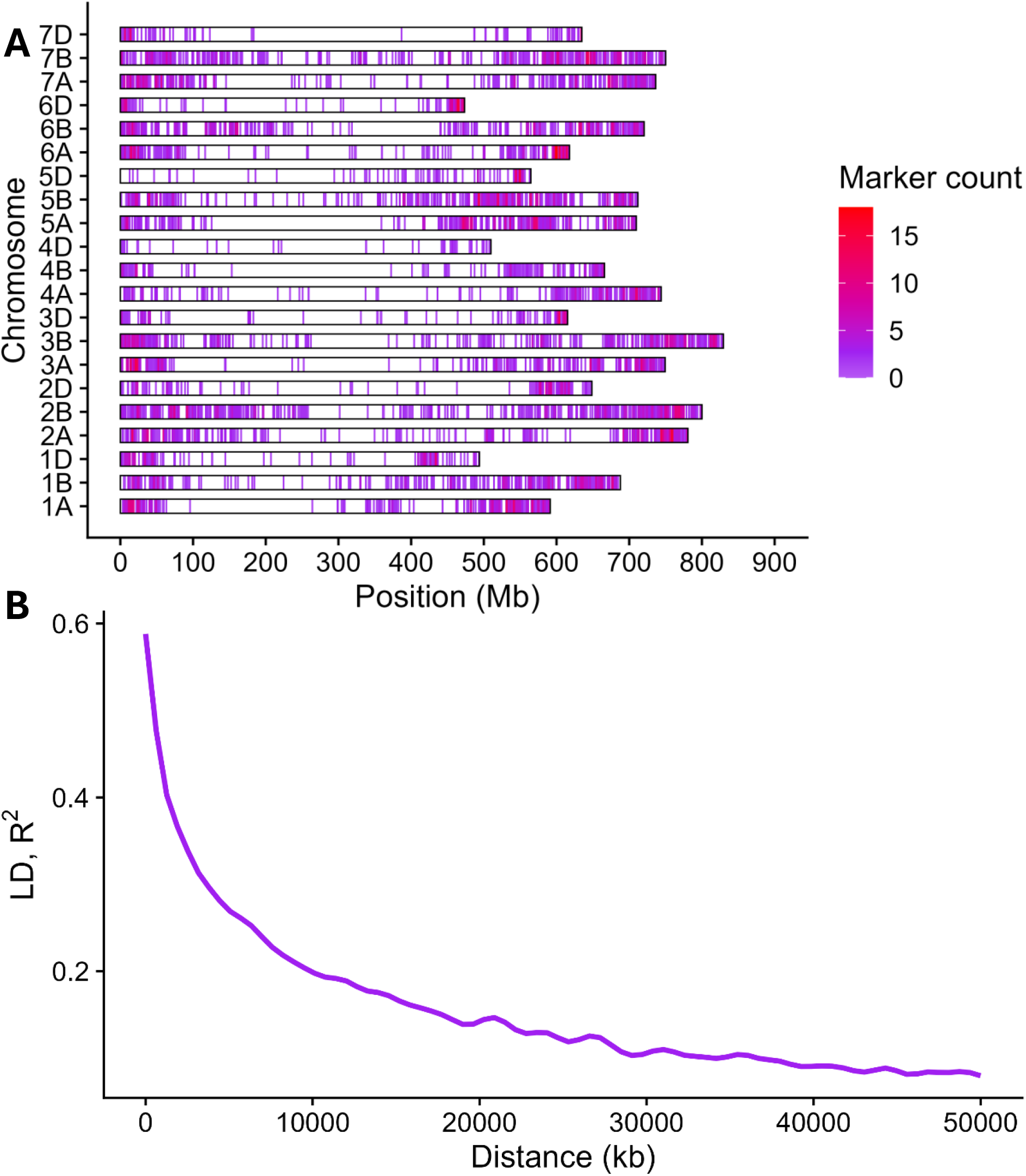
A. Distribution of the 6054 markers across the chromosomes in 2.5 megabase (Mb) windows. B. Linkage disequilibrium (LD) decay (kilobases, kb). Decay line was computed using a generalized additive model.

**Supplementary Figure 5.**
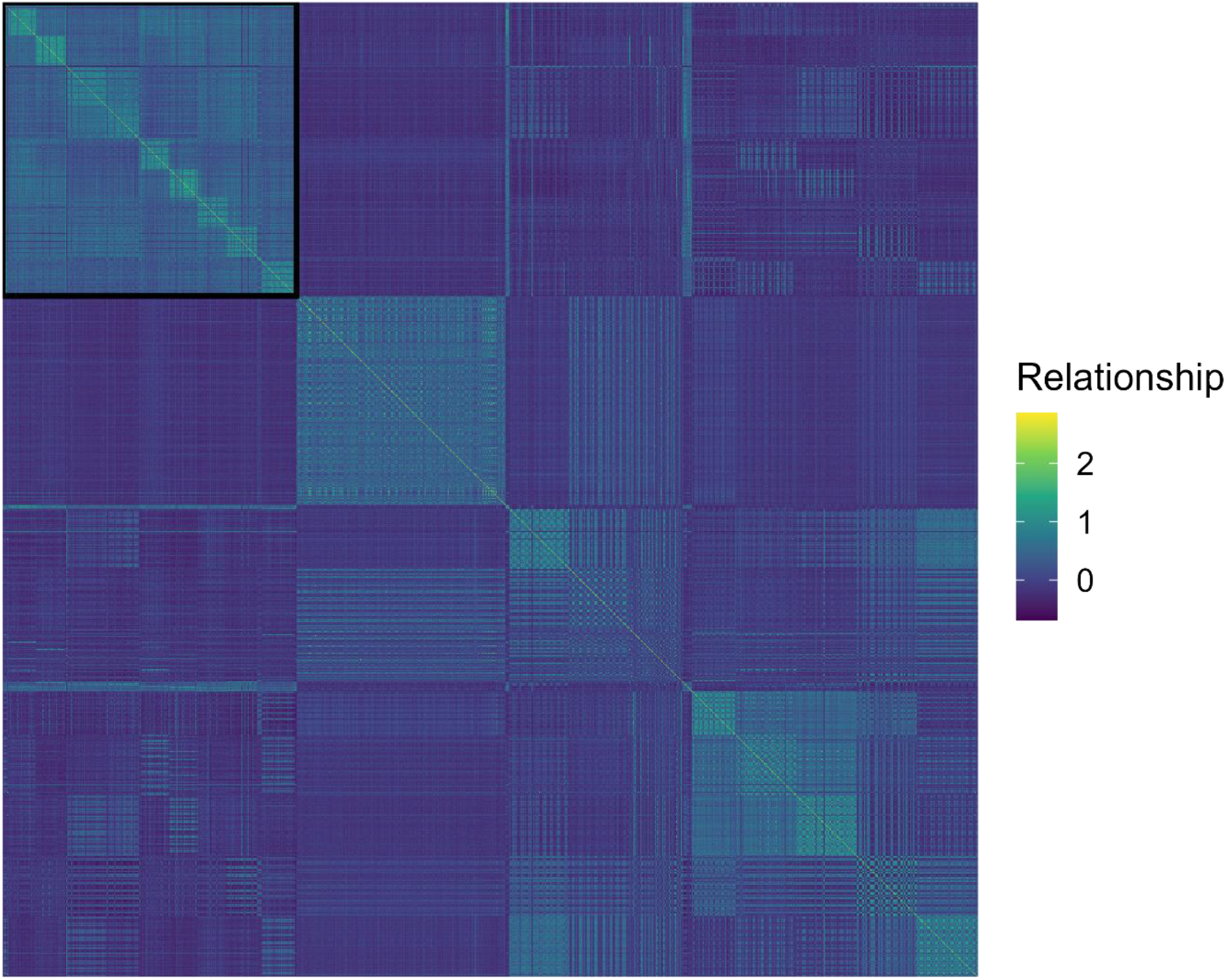
Heatmap of the genomic relationship matrix based on Van Raden’s method 1 (VanRaden, 2008) of all 1474 lines. The Suntop-derived nested association mapping lines are boxed in the upper-left corner.

## References

Amin, A., Christopher, J., Cooper, M., Collins, B., Voss-Fels, K., Hickey, L., and Chenu, K. (2025). Envirotyping facilitates understanding of genotype× environment interactions and highlights the potential of stay-green traits in wheat. Field Crops Res. 331:109940.

Bourdon, R. M. (2000). Understanding animal breeding. Prentice Hall Upper Saddle River, NJ.

Butler, D. G., Cullis, B. R., Gilmour, A. R., Gogel, B. J., and Thompson, R. al (2017). ASReml-R reference manual version 4. VSN Int. Ltd Hemel Hempstead HP1 1ES UK Advance Access published 2017.

Chang, C. C., Chow, C. C., Tellier, L. C., Vattikuti, S., Purcell, S. M., and Lee, J. J. (2015). Second-generation PLINK: rising to the challenge of larger and richer datasets. Gigascience 4:s13742–015-0047–8.

Cole, J. B., and VanRaden, P. M. (2011). Use of haplotypes to estimate Mendelian sampling effects and selection limits. J. Anim. Breed. Genet. 128:446–455.

Cooper, M., and Messina, C. D. (2023). Breeding crops for drought-affected environments and improved climate resilience. Plant Cell 35:162–186.

Daetwyler, H. D., Hayden, M. J., Spangenberg, G. C., and Hayes, B. J. (2015). Selection on optimal haploid value increases genetic gain and preserves more genetic diversity relative to genomic selection. Genetics 200:1341–1348.

Endelman, J. B. (2011). Ridge regression and other kernels for genomic selection with R package rrBLUP. Plant Genome 4.

Falconer, D. S., Mackay, T. F., and Frankham, R. (1996). Introduction to quantitative genetics (4th edn). Trends Genet. 12:280.

Fan, B., Onteru, S. K., Du, Z.-Q., Garrick, D. J., Stalder, K. J., and Rothschild, M. F. (2011). Genome-Wide Association Study Identifies Loci for Body Composition and Structural Soundness Traits in Pigs. PLoS ONE 6:e14726.

Goiffon, M., Kusmec, A., Wang, L., Hu, G., and Schnable, P. S. (2017). Improving response in genomic selection with a population-based selection strategy: optimal population value selection. Genetics 206:1675–1682.

Habier, D., Fernando, R. L., and Dekkers, J. C. (2009). Genomic selection using low-density marker panels. Genetics 182:343–353.

Hayes, B. J., Mahony, T. J., Villiers, K., Warburton, C., Kemper, K. E., Dinglasan, E., Robinson, H., Powell, O., Voss-Fels, K., and Godwin, I. D. (2024). Potential approaches to create ultimate genotypes in crops and livestock. Nat. Genet. 56:2310–2317.

Heffner, E. L., Sorrells, M. E., and Jannink, J. (2009). Genomic Selection for Crop Improvement. Crop Sci. 49:1–12.

Henderson, C. R. (1975). Best linear unbiased estimation and prediction under a selection model. Biometrics Advance Access published 1975.

Hunt, J. R., Lilley, J. M., Trevaskis, B., Flohr, B. M., Peake, A., Fletcher, A., Zwart, A. B., Gobbett, D., and Kirkegaard, J. A. (2019). Early sowing systems can boost Australian wheat yields despite recent climate change. Nat. Clim. Change 9:244–247.

Kemper, K. E., Bowman, P. J., Pryce, J. E., Hayes, B. J., and Goddard, M. E. (2012). Long-term selection strategies for complex traits using high-density genetic markers. J. Dairy Sci. 95:4646–4656.

Liu, H., Zhou, H., Wu, Y., Li, X., Zhao, J., Zuo, T., Zhang, X., Zhang, Y., Liu, S., and Shen, Y. (2015). The impact of genetic relationship and linkage disequilibrium on genomic selection. PloS One 10:e0132379.

Lloyd, A. (2023). Crossover patterning in plants. Plant Reprod. 36:55–72.

Lush, J. L. (1937). Animal Breeding Plans. Collegiate Press, Incorporated.

Meuwissen, T. H. E., Hayes, B. J., and Goddard, M. E. (2001). Prediction of Total Genetic Value Using Genome-Wide Dense Marker Maps. Genetics 157:1819–1829.

Piepho, H. P., Möhring, J., Melchinger, A. E., and Büchse, A. (2008). BLUP for phenotypic selection in plant breeding and variety testing. Euphytica 161:209–228.

Quitzow, R., Balmaceda, M., and Goldthau, A. (2025). The nexus of geopolitics, decarbonization, and food security gives rise to distinct challenges across fertilizer supply chains. One Earth 8.

Ray, D. K., Gerber, J. S., MacDonald, G. K., and West, P. C. (2015). Climate variation explains a third of global crop yield variability. Nat Commun 6: 5989.

Scrucca, L. (2016). On some extensions to GA package: hybrid optimisation, parallelisation and islands evolution. ArXiv Prepr. ArXiv160501931 Advance Access published 2016.

Shaffer, W., Papin, V., Yadav, S., Voss-Fels, K. P., Hickey, L. T., Hayes, B. J., and Dinglasan, E. G. (2025). Local genomic estimates provide a powerful framework for haplotype discovery. bioRxiv Advance Access published 2025.

Snowdon, R. J., Wittkop, B., Chen, T.-W., and Stahl, A. (2021). Crop adaptation to climate change as a consequence of long-term breeding. Theor. Appl. Genet. 134:1613–1623.

Solberg, T. R., Sonesson, A. K., Woolliams, J. A., and Meuwissen, T. (2008). Genomic selection using different marker types and densities. J. Anim. Sci. 86:2447–2454.

Tong, J., Tarekegn, Z. T., Jambuthenne, D., Robinson, H., Pandit, M., Villiers, K., Periyannan, S., Hickey, L., Dinglasan, E., and Hayes, B. J. (2025). Haplotype stacking to improve stability of stripe rust resistance in wheat. Theor. Appl. Genet. 138:267.

VanRaden, P. M. (2008). Efficient methods to compute genomic predictions. J. Dairy Sci. 91:4414–4423.

Villiers, K., Dinglasan, E., Hayes, B. J., and Voss-Fels, K. P. (2022). genomicSimulation: fast R functions for stochastic simulation of breeding programs. G3 12:jkac216.

Villiers, K., Voss-Fels, K. P., Dinglasan, E., Jacobs, B., Hickey, L., and Hayes, B. J. (2024). Evolutionary computing to assemble standing genetic diversity and achieve long-term genetic gain. Plant Genome 17:e20467.

Voss-Fels, K. P., Stahl, A., Wittkop, B., Lichthardt, C., Nagler, S., Rose, T., Chen, T.-W., Zetzsche, H., Seddig, S., and Majid Baig, M. (2019). Breeding improves wheat productivity under contrasting agrochemical input levels. Nat. Plants 5:706–714.

Wang, Z., Miao, L., Tan, K., Guo, W., Xin, B., Appels, R., Jia, J., Lai, J., Lu, F., and Ni, Z. (2025). Near-complete assembly and comprehensive annotation of the wheat Chinese Spring genome. Mol. Plant 18:892–907.

Wray, N. R., and Goddard, M. E. (1994). Increasing long-term response to selection. Genet. Sel. Evol. 26:431–451.

Yadav, S., Dillon, S., McNeil, M., Dinglasan, E., Mago, R., Dodds, P., Hickey, L., and Hayes, B. J. (2025). Optimising parent selection in plant breeding: comparing metaheuristic algorithms for genotype building. Theor. Appl. Genet. 138:242.

